# Emergence of a synergistic scaffold in the brains of human infants

**DOI:** 10.1101/2024.02.23.581375

**Authors:** Thomas F. Varley, Olaf Sporns, Nathan J. Stevenson, Martha G. Welch, Michael M. Myers, Sampsa Vanhatalo, Anton Tokariev

## Abstract

The human brain is a complex organ comprising billions of interconnected neurons which enables interaction with both physical and social environments. Neural dynamics of the whole brain go far beyond just the sum of its individual elements; a property known as “synergy”. Previously it has been shown that synergy is crucial for many complex brain functions and cognition, however, it remains unknown how and when the large number of discrete neurons evolve into the unified system able to support synergistic interactions. Here we analysed high-density electroencephalography data from late fetal to early postnatal period. We found that the human brain transitions from redundancy-dominated to synergy-dominated system around birth. Frontal regions lead the emergence of a synergistic scaffold comprised of overlapping subsystems, while the integration of sensory areas developed gradually, from occipital to central regions. Strikingly, early developmental trajectories of brain synergy were modulated by environmental enrichment associated with enhanced mother-infant interactions, and the level of synergy near term equivalent age was associated with later neurocognitive development.

## 1 Introduction

The human brain is a paradigmatic example of a complex system displaying emergent structure. Despite being composed of billions of individually functioning neurons, their collective dynamics leads to the emergence of a unified “whole” capable of integrating information, learning, and surviving in complex environments. The property where a system collectively shows structure that is irreducible to the sum of its parts is known as synergy [1], and it is thought to play a key role in the self-organization of the brain into a unified whole [2, 3]. Understanding this process is a fundamental challenge in modern neuroscience. Information theory has emerged as a core tool kit for the analysis of modern complex systems science [4], and has been used to explore the structure of information processing and cognition in the brain at multiple scales [5–7]. Previous fMRI studies on adults have shown that synergistic sub-systems are widespread across the cortex [7, 8], and that the distribution of synergies changes across the adult lifespan [5].

In the development of an individual’s brain, functional synergy can only emerge after the development of underlying structural brain networks. It is currently well established that most structural and functional organization in brain networks takes place during the few months around birth [9–14]. This process is driven by a combination of genetically guided growth of the major structural networks and an activity-dependent organization of the functional networks [15–18] into integrated and segregated ensembles that together form the synergistic whole [2, 19]. However, it is not known how and when synergistic brain function appears, and whether it emerges sequentially or uniformly across the newly developed cortex. Spontaneous cortical activity in large neuronal ensembles directly facilitates the emergence of the brain’s functional synergistic structure. Recording this activity provides a natural test-bed to study the self-organization of higher-order dependencies in the brain. In this paper, we use information theory to assess the emergence of a “synergistic scaffold” in the functional architecture brain during late fetal and early postnatal development.

We hypothesized that synergy emerges in spatially resolved sequences during the early development of the brain, and moreover, that the emergence of a consolidated “synergistic scaffold” links to later neurocognitive performance at individual level. To test this, we used a recently proposed measure of higher-order structure in complex systems: the O-Information (Ω) [20]. The O-information can assess whether statistical structure of a complex system, such as global EEG activity, is synergy-dominated (i.e., the collective whole contains information that is irreducible to smaller collections of parts) or redundancy-dominated (i.e., the collective whole can be compressed or simplified by pruning duplicated information). For a given multivariate system **X**, if Ω(**X**) *<* 0, then the system is synergy dominated, and if Ω(**X**) *>* 0, then the system is redundancy dominated. In this study, we determined the level and topographic distribution of O-information during early development of spontaneous cortical activity in newborn infants. First, we estimated the O-information across the whole brain to assess global information structure in the EEG. Second, we extracted maximally synergistic subsystems to identify regional subsets with highest synergy. This yielded spatially resolved tracking of the emerging synergistic subsystems in fine-grained detail, which could then be compared to functional hierarchies and later neurocognitive development.

## 2 Results

### 2.1 Synergy in the brain increases during early maturation

We found that O-information becomes increasingly negative with age at all studied frequency bands (Figure 1A; *ρ < −*0.42, pFDR *<* 0.001, Spearman test), indicating a transition from a redundancy-dominated to a synergy-dominated structure. The effect size was the strongest at higher frequencies (alpha, 8-13 Hz, *ρ* = *−*0.57; and beta, 13-22 Hz, *ρ* = *−*0.59). These preterm infants, when assessed at postconceptional ages less than term age, showed high positive O-information values suggesting a prevalence of redundancy-dominated dynamics, while the O-information levels shifted towards zero and even became negative around term age. Once again reflecting increasing presence of synergy, and the transformation of the brain into a more synergy-dominated system. This is consistent with the works demonstrating changes in connectivity networks in the preterm brain during early development [21, 22]. Assessing the strength (O-information) of the most synergistic ensembles of brain regions, across all possible subsystem sizes, in individual infants (Figure 1B, top row) revealed systematic increases in negative O-information across all temporal scales as a function of age. At the same time, the size of the optimal system (or “synergistic subsystem”) grew at an accelerated pace near term age (Fig.1B, bottom row). We also found that expansion of the “synergistic subsystem” across the whole cortex and increase of the overall synergy are two highly correlated signatures of early brain development (Figure 1C; *ρ < −*0.65, pFDR *<* 0.001 for all, Spearman test).

**Fig. 1.**
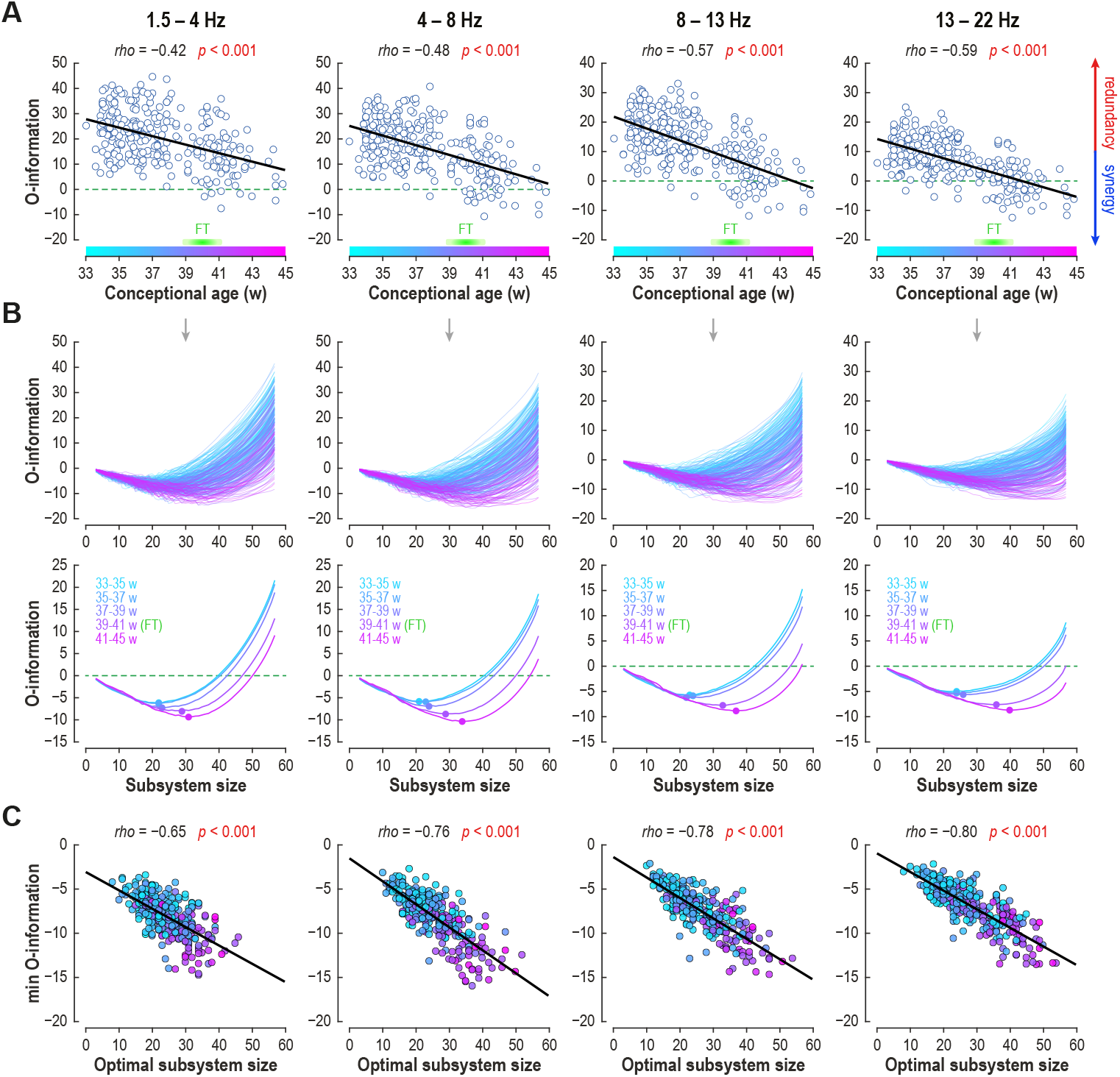
Emergence of synergistic brain during the three months around birth. **A**, The global values of the O-information becomes significantly more negative at all inspected oscillatory frequencies (Spearman) pointing to emerging synergy towards full term (FT) age. **B**, Relationship between optimally synergistic O-information and the subsystem size in individual infants (top row) and in averaged age groups (bottom row), coloured for the infant’s age at EEG recording. The dots in the bottom plots depict the minimum of O-information curves that reflects the size of subsystems with maximum synergy in the given age group (the optimal “synergistic subsystem”). Note, the systematic growth of the synergistic subsystem with age at all frequencies, with particularly rapid changes around term age. **C**, Relationship (Spearman) between O-information levels and the size of the maximally synergistic subsystem in individual EEG (computed from the curves on the top row in B; colours code the age at recording). Developmental expansion of the synergistic subsystem and increase of its synergistic capacity are highly correlated (the brain becomes more complex).

### 2.2 The synergistic scaffold expands sequentially from the frontal to other brain regions

Next, we analysed the early development of the spatial configuration of the “synergistic scaffold” by comparing node consistency maps in three age groups: early preterm, late preterm, and early postnatal. These maps indicate participation frequency of the given cortical region (or “node”) in the synergistic scaffold (Figure 2). In the early preterm infants, the synergistic scaffold was dominated by a symmetric frontal cluster, and a markedly less prominent occipital cluster, that was more clearly expressed at higher (alpha and beta; 8-22 Hz) frequencies (Figure 2, top row). In comparison, the late preterm group showed spatial expansion and an increase in consistency across subjects for the frontal and occipital clusters in a whole frequency range (Figure 2, middle row). In the early postnatal age group, the synergistic scaffold had expanded considerably to include the central cortical areas as well (Figure 2, bottom row). This sequential recruitment of cortical areas is consistent with recent studies in adults that found that high-synergy systems tended to straddle multiple canonical functional networks [7, 19].

**Fig. 2.**
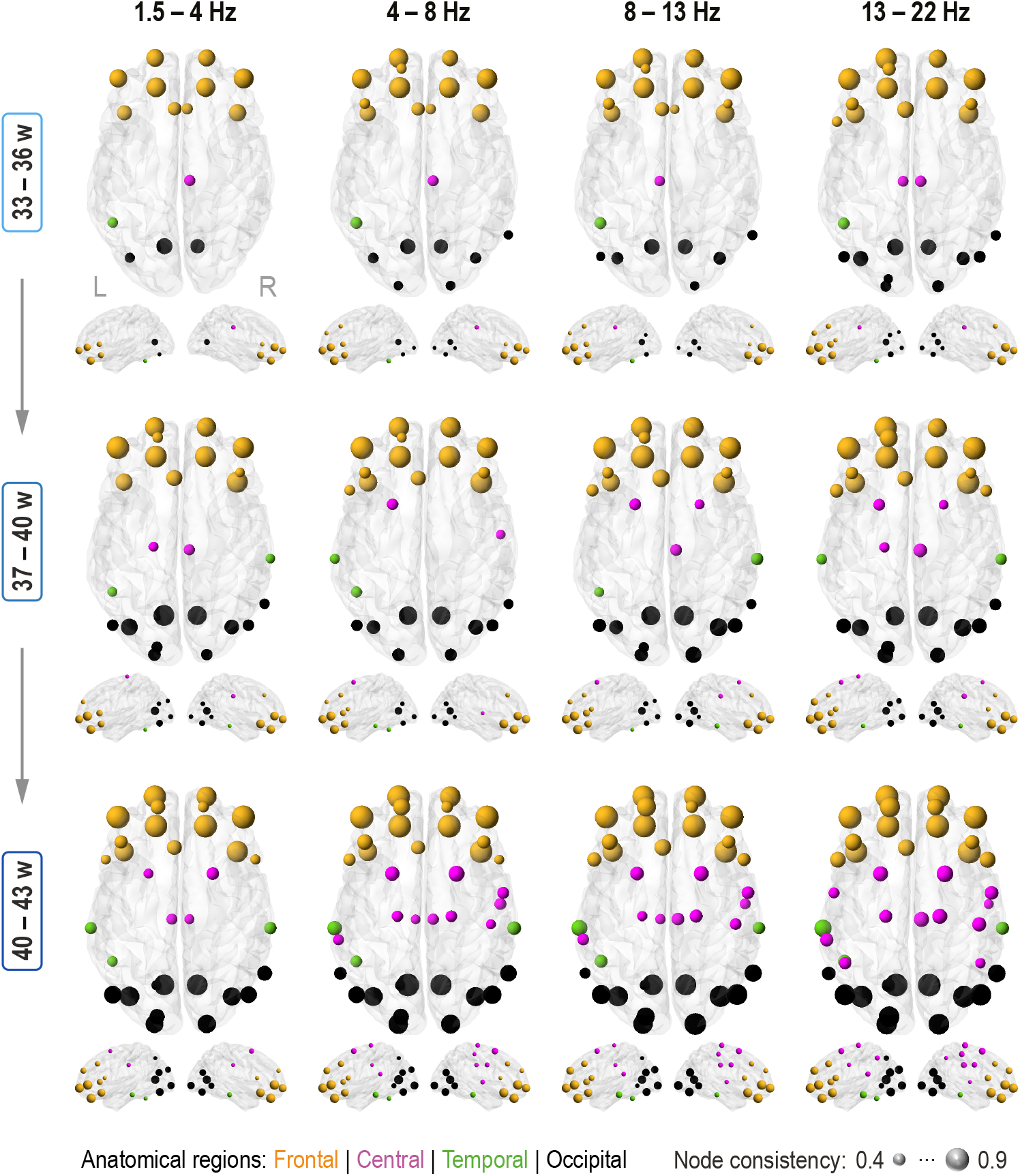
Early spatial development of the synergistic scaffold at different frequency bands. The ball size indicates participation frequency, of each cortical region (“node”) in the optimally synergistic ensembles (collectively forming the “synergistic scaffold”) in different age groups (rows), and at different frequency bands (columns). For each infant, maximally synergistic subsets of the whole brain were discovered using simulated annealing [23], with the negative O-information as the objective function (for details see Materials and Methods). Running the optimizer a large number of times reveals a landscape of non-identical, but overlapping, synergistic ensembles, which recruit brain regions from different areas at different periods of development, so we extracted the number of times each node was selected across all trials of all infants, and visualize [24] those that appear in *>* 40% of optimal sets. Note the early prominence of frontal synergistic scaffold, the incremental recruitment of the occipital regions, and the late-appearing participation of the central regions. The colours indicate the anatomical affiliation in different cortical regions: frontal (orange), central (purple), temporal (green), and occipital (black).

### 2.3 Newborn brain synergy precedes long-term neurocognitive development

We then asked if the level of synergy at the time of normal birth has neurodevelopmental implications. To this end, we correlated O-information metrics taken around term age (38-42 weeks) with later neurocognitive performance assessed at 18 months using standardized Bayley Scales (available for *N* = 41 subjects; [25, 26]). The later neurocognition was robustly correlated to the individual O-information levels across the whole frequency range of interest (Figure 3A; *ρ ≤ −*0.41, pFDR *≤* 0.008). These correlations were not affected after regressing age at EEG recording from the O-information values (see bottom row on Figure 3A). Other characteristics of synergistic subsystems were also linked to later neurocognitive development. The minimum O-information showed negative correlation with later performance (Figure 3B, top row; *ρ ≤ −*0.32, pFDR *≤* 0.039), suggesting that the early emergence of synergistic structures may promote cognitive development. Similarly, the size of the synergistic subsystem was strongly correlated with better cognitive performance (Figure 3B, bottom row; *ρ ≥* 0.46, pFDR *≤* 0.002), suggesting that later cognitive development may benefit from including wider brain areas in the synergistic whole. Taken together, our results suggest that the level of overall synergistic cortical activity as well as its spatial expansion over the cortex strongly reflect an individual’s capacity for neurocognitive development.

**Fig. 3.**
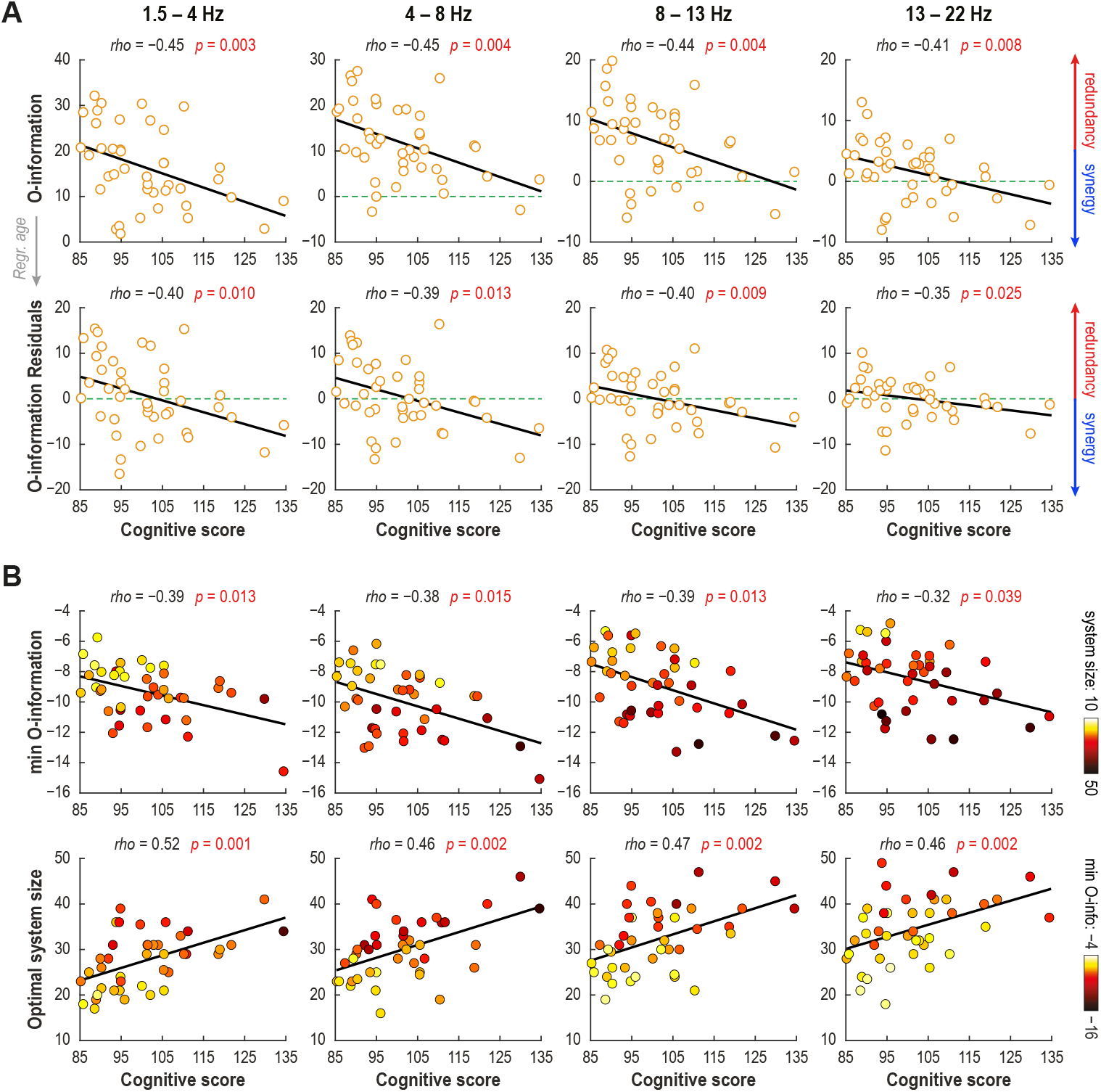
Synergy-dominated organization of the maturing brain associates with better neurodevelopment. **A**, Correlation of the global O-information in the newborn brain and cognitive performance at 18 months of age. The upper panel shows the original values, whereas the lower one presents values after regressing age at EEG. **B**, Both properties of the optimal sub-system: minimum O-information (upper row) and optimal sub-system size (lower row) correlate with future cognitive development. Colours of the dots cross-link two properties. Overall, larger synergistic structures, with higher level of synergy (more negative values of O-information minima), together were associated with better outcomes across all frequency bands. All EEG measures were taken from recordings near term age (38-42 weeks). The correlation analyses were done using Spearman test, and significance estimates are corrected post-hoc for multiple comparisons.

### 2.4 Environmental enrichment modulates early developmental trajectories of brain synergy

Intertwined with the genetically programmed schedules of brain development [27, 28], there is growing evidence that early activity-dependent brain development can be substantially affected by environmental factors [16, 18, 29–32]. These “acquired effects” range from major medical adversities like preterm birth or perinatal asphyxia [33–36] to more subtle issues [31, 37–39]. Recent preclinical and clinical studies have suggested that various changes in an infants’ living environment (“environmental enrichments”) may support improved neurodevelopment at many levels of inspection from the cellular level [40] to brain networks [22, 41, 42], brain structure [43], and many aspects of later neurobehavioural outcomes [26, 44]. Therefore, we assessed whether the early emergence of synergistic brain activity could be affected by the living environment.

To this end, we took advantage of having two subgroups in our dataset: half of the infants had been treated according to all the evidence-based guidelines of preterm care (standard care group, SC); the other half was assigned to a group that received additional intervention during their stay in the neonatal intensive care unit to facilitate emotional parent-infant connection (Family Nurture Intervention, FNI; [45], which is considered to be biologically relevant, and medically mild addition to nursing practice [45–48].

The developmental trajectories of O-information using two-weeks-wide time windows were clearly different between these subgroups (Wilcoxon rank-sum test; Figure 4A). The SC infants showed a bi-phasic trajectory with an initial plateau until near term age (*≈*38 weeks) followed by an abrupt transition towards greater synergy, whereas the FNI infants showed a steady increase in synergy in O-information. The groups were consistently different across all frequencies at two time points: 35 weeks (FNI *>* SC; *p ≤* 0.051 for all); and 38 weeks (FNI *<* SC; *p ≤* 0.033 for all) with the strongest effect in the theta band (4-8 Hz; *p* = 0.007, effect size *r* = 0.49). The neighbouring age bins showed concordant though less significant differences (FNI *>* SC at week 34 for 4-13 Hz and FNI *<* SC at weeks 39-40 for 4-22 Hz; *p <* 0.1 for all). The Oinformation minima (Figure 4B) was lower in the SC infants (*p <* 0.1) at 35 weeks in theta (4-8 Hz) and beta (13-22 Hz) bands, whereas at 38 weeks it became significantly more negative for FNI (4-22Hz; *p ≤* 0.07, effect size 0.32 *< r <* 0.46). In turn, the optimal system size (Figure 4C) at 35 weeks was larger in the SC group across the full bandwidth (*p <* 0.1), and most prominently in the beta band (13-22 Hz; *p* = 0.008, effect size *r* = 0.32), but at 38 weeks it changed to opposite (*p <* 0.034, effect size 0.38 *< r <* 0.44) except in the alpha band (8-13 Hz). Overall, these findings suggest that the synergistic scaffold in the brains of FNI infants develops more dynamically, yet steadily, before reaching term age. The cumulative effect of environmental enrichment intervention more likely causes the most prominent group difference just prior normal birth.

**Fig. 4.**
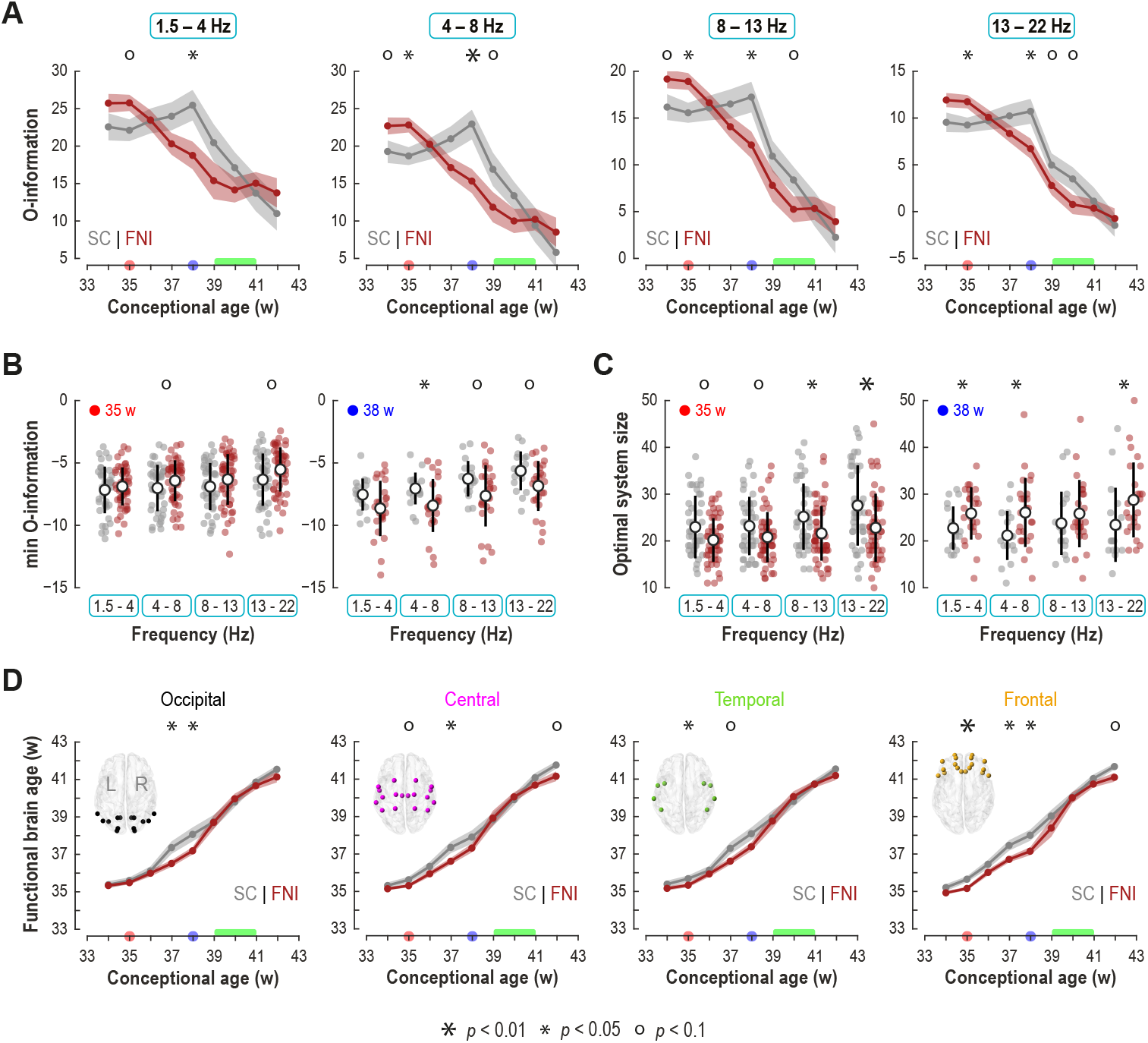
Environmental enrichment modulates global synergy and local maturity of neuronal activity. **A**, Developmental trajectories in the global synergy (O-information) in the SC (grey) and FNI (dark red) groups at different frequency bands. Dots indicate group mean values computed at bi-weekly intervals, with shades depicting standard error of the mean (SEM); asterisks and circles indicate significant group differences (Wilcoxon rank-sum test, uncorrected). The red and blue dots on the X-axis depict age points with the strongest consistent difference across all frequencies, which were selected for further analysis (in B). The green bar depicts period corresponding to normal full-term birth. Note the differences in developmental trajectories: FNI shows steady decrease, whereas SC is characterized by an initial “plateau” until a rapid decline occurs after about 38 weeks of age to reach the levels comparable to the FNI group. **B**, Comparison of SC and FNI groups for Oinformation minima across different frequencies at the two time points marked in A (left, 35 weeks; right, 38 weeks). **C**, Comparison of the optimal synergistic subsystem size for the same time points as in B. In both B and C, individuals’ values are shown with dots and the whiskers show mean (circle) and standard deviation (thick line), respectively. **D**, Regional trajectories of local maturation in neuronal activity estimated by functional brain age (FBA) in the SC (grey) and FNI (red) groups. The FBA values (dots show infant group means; shade depicts SEM) in the graphs were computed as average of parcels in each cortical region.

We then wanted to understand whether the environmental effects on the synergy development is reflected at the level of local maturation of neuronal activity, as assessed by independent, machine learning-based estimates of the functional brain age (FBA [49]) at each cortical parcel. There were significant, region-specific group differences (Wilcoxon rank-sum test) in the maturation of cortical neuronal activity (Figure 4D). The group difference (SC *>* FNI) was first seen in the frontal regions (35 weeks; *p* = 0.007); it expanded to the whole cortex before term age (37 weeks; *p <* 0.07), followed by frontal and occipital group differences at around term age (38 weeks; *p <* 0.05). At term age, groups did not differ until week 42 when SC infants showed higher FBA in frontal and central regions (*p <* 0.1).

Taken together, the environmental enrichment appears to cause co-directional changes in the developmental trajectories of both the local neuronal activity (FBA) and the system-wide synergy (O-information), with the most prominent manifestation in the frontal regions (see also Figure 2).

## 3 Discussion

Our results indicate that system-level organization of the infant brain is characterized by an early, region-specific shifting from a functional structure dominated by redundant interactions to one dominated by synergistic interactions. Synergistic structure first emerges in the frontal lobe before spreading over other cortical areas in a specific spatiotemporal sequence, and this developmental trajectory can be modified by simple environmental modulations. The neurodevelopmental implication of redundancy/synergy balance was demonstrated by its significant correlation to later emerging neurocognitive functions.

Prior work has established the presence of synergy in human fMRI data [2, 7, 19], and that the distribution of redundancies and synergies can change gradually during aging [5], or more rapidly due to changes in level of consciousness [2, 3]. However, it has remained unclear to what extent these higher order synergies, beyond their statistical significance, influence the mechanisms of behaviour and cognition. Our results show that infant synergy is significantly associated with later cognitive performance. This finding suggests that the ability of the brain to generate and maintain higherorder synergies at earlier stages facilitates the development of complex cognition later in life. This is consistent with work on artificial neural networks, where the synergy in individual “neurons” facilitates the capacity of the system to engage in multitasking and other integrative behaviours [50]. Similarly, the finding that environmental enrichment early in life modulates the development of the synergistic structure suggests that the presence (or absence) of synergy is at least partially mediated by complex multisensory experiences.

Synergy may provide a fundamental mechanism that supports complex, higher brain functions characteristic of human behaviour. Many higher neurobehavioural abilities in humans need efficient synergy to support communication within and between brain regions. The cellular underpinnings are characterized at all levels of neuronal communication, from synapses and cell types to different spatial scales in neuronal networks [51]. In the context of the present findings, neuronal networks are likely the most fruitful level of explanation: At the microscale, comparisons to species that are phylogenetically more distant, such as rodents, indicate that the vast expansion of the human cortex comes with an evolutionary emergence of much denser interneuronal circuitry [52] to facilitate local information processing. At the mesoscale level, pyramidal neurons in the human cortex have evolved exhibiting increased synaptic connectivity and many other unique input–output integration properties [51, 53], which together improve information processing across cortico-cortical circuitries.

At the macroscale level, which is directly comparable to our present results, evolution of cortico-cortical network organization [54, 55] appears to be characterized by tuning the quality more than the quantity of connections. Despite larger brain size and higher anatomic variability in the human brain [56], there appears to be a wider optimization of connection strengths (“weights”) that supports efficient macroscale information transfer [57, 58]. The early growth and organization of human neuronal networks is characterized by a prominent temporal overlap and prolongation of phases that would be temporally far more distinct in most other species [51, 56, 59]. Such gradual network development provides an ideal framework for an activity- and experience-dependent development of macroscale synergy, i.e. a change in the functional network characteristics that needs rapid optimization of the connection strengths in the newly developed cortico-cortical networks. Moreover, this rationale would offer a mechanistic explanation for the finding that environmental enrichment may modulate the trajectories in synergy development: While the initial growth of neuronal connections is supported by the genetic code and endogenous neural activity [15, 18, 28], the following global organization including synergy in the neuronal networks is guided by neural activity that is sensitive to environmental and other acquired effects [16, 34, 35, 44]. The frontal lead in the emergence of functional synergy maybe somewhat counterintuitive with respect to the general frontal delay in brain maturation [60–63]. However, the existing literature is mainly based on structural measures whereas synergy in network activity characterizes informational signatures of macroscale interaction between neuronal ensembles: changes in the frontal region here reflect changes in the interactions between frontal neurons and potentially the rest of the brain. Recent studies have, indeed, highlighted the developmentally and functionally hierarchical brain organization, with clear gradients between sensory and association areas [64–67]. The sensory areas develop to process information from inputs specific to the respective sensory modality that are only later communicated to higher order systems during postnatal life. In contrasts, development of association areas, especially the frontal regions, is largely characterized by optimizing information processing in the global cortico-cortical networks. Thus, emergence of macroscale synergy is inherently linked to early development of cortico-cortical circuitries in association areas, whereas corresponding synergy emerges much later in the sensory cortical areas. The frontal lead in synergy development is also intriguingly compatible with our observation that the trajectories of synergy development are modulated by environmental enrichments (FNI group). Recent studies have established that human brain shows particularly protracted developmental time spans across many levels (Wallace and Pollen, 2024): The anatomical studies from the months around birth indicate several months-long co-existence of overlapping macroscale connectivity [68], whereas many synaptic characteristics [60, 69, 70] or molecular expression profiles [69, 71] show a strong neoteny persisting until late childhood. Such a protracted and gradual development renders the system modifiable, or able to learn, through the well-established activity-dependent process; however, it also makes the developing brain subject to environmental effects, such as perturbations by medical adversities. Here, we postulate that the trajectory of synergy development in the FNI group represents the natural course as it would happen in utero. Conversely, the biphasic trajectory in the SC group reflects an initial slowing down of synergy development by the prematurity-related medical adversities [34], followed by a rapid catch-up near-term age. Intriguingly, the transient developmental FNI effect was also seen in the independent maturation measure (FBA) of the local neuronal activity. Prior work on the newborn corticocortical networks showed that prematurity leads to persistent changes [33, 72–75], which may be substantially reduced by environmental enrichment [22]. Our present results and prior studies together suggest that environmental effects may substantially modulate the early organization of functional brain networks. Both the early development of cortico-cortical network activity and the emergence of brain synergy build on co-stabilization of the long-range axonal conduction via myelination [76, 77] and the local synaptic transmission [70]. Experimental studies have indicated these factors in recovery from brain injury [78, 79], learning and development [77], as well as in the brain response to early environmental enrichment interventions [80, 81].

While it appears widely accepted that very early environmental modulation, or nurturing, can improve later neurodevelopment [82], understanding the underlying neuronal mechanisms needs substantially more systematic and translational work, preferably in phylogenetically aligned animal models.

Finally, it is worth reflecting on the implications of these, and other, results that use higher-order information theory to explore brain activity. The standard approaches based on single regions, or pairwise region-to-region interactions (commonly used in network models) capture only a very small subset of all possible dependencies that co-exist and interact in the brain. Our present approach with a very dense scalp EEG recording allowed assessment of approximately 3 thousand possible directed interactions, which is at the physical limits of spatial resolution available in studying neuronal activity in human infants [33]. If we consider the possibility of higher-order synergies up to the 5^th^ order, then the total number of possible dependencies expands to approximately 7 million interactions. The network then represents only 0.0005% of the possible structure of the system (stopping at 5^th^ order interactions; we can, of course, go higher). This “shadow structure” [7, 19] represents a vast space of largely unexplored structured brain activity. Our results here show that this space contains patterns of multivariate information that are associated with, and possibly promote, specific aspects of human development, cognition, and parent-child interactions. Future work exploring this higher-order space may yet yield further, fruitful insights into the nature of brain, mind, and behaviour.

## 4 Methods

### 4.1 Subjects and background information

We analysed EEG dataset from N = 135 preterm infants (born at 31.1 *±* 2.4 weeks; mean *±* std) that was collected during the Family Nurture Intervention (FNI) randomized controlled trial (#NCT01439269 in ClinicalTrials.gov) in Morgan Stanley Children’s Hospital of New York at the Columbia University Medical Center [45]. The general dataset consisted of two subgroups: standard care infants (SC; *N* = 61, born 30.9 *±* 2.5 weeks), and those who additionally underwent intervention aimed to facilitate mother-infant emotional connection (FNI; *N* = 74, born 31.2 *±* 2.3 weeks). Subgroups showed no difference in the age of birth (*p* = 0.76, Wilcoxon rank sum test). Review Board at Columbia University Medical Center (NY, USA) approved study and recruitment procedures. Written consents were obtained from mothers before the start of the intervention.

### 4.2 EEG recordings

EEG data was collected longitudinally between weeks 33 and 45 of conceptual age during daytime sleep using 128-channel system Electrical Geodesics system. Each session lasted for about one hour and covered at least one full sleep cycle comprising periods of active and quiet sleep [83]. Four facial electrodes were excluded from further analysis leading to final 124 EEG channels per infant. During the recordings the impedance of EEG electrodes was kept below 50 kiloohms. Original recordings were done using vertex electrode as a reference, with a sampling rate Fs = 1 kHz, and using band-pass filter 0.1–400 Hz. After the recording, all data were re-referenced to average montage.

### 4.3 EEG pre-processing

First, we identified periods of stable quiet sleep (precursor of non-rapid eye movement sleep) from the whole recording using conventional criteria [84]. We used quiet sleep because EEG during this state is phenomenologically more discriminative: it has discontinuous structure comprising bursts of activity and in-between silent intervals [73, 85]. Moreover, quiet sleep EEG epochs are technically more stable: they contain less electromyographic and electrooculographic artifacts as well as noise associated with movement. Next, all selected epochs were visually reviewed to identify prominent artifacts such as bad or absence of skin-electrode contact, presence of electromyogram, and cable movement. Further we applied independent component analysis (ICA) to check for the presence of electrocardiographic artifact in every recording and removed it where it was detected. ICA was also used to check and clean rare artifacts caused by interference of other medical devices in neonatal intensive care unit. For the analysis we selected 5 minutes of artifact-free EEG by combining ten equidistant 30-second-long windows across the whole recording. That was done to overcome variability in available EEG lengths across different subjects and to obtain representative epochs that characterize the whole period of quiet sleep. Channels that were bad across entire recording session were removed from further analysis. The final dataset included N = 289 EEG recordings (from N = 134 subjects) which satisfied quality and length requirements: N = 30 infants with one recording, N = 51 with two recordings, N = 51 with three recordings, and N = 1 with four recordings. These recordings then were band-pass filtered within frequency range 0.4–40 Hz and down sampled to Fs = 100 Hz.

### 4.4 Computation of cortical signals

Scalp-level EEG signals were reconstructed into cortical signals using three-shell general infant head model [9, 86]. The model included scalp, skull, and intracranial volume boundaries approximated with 2562 vertices per compartment and having conductivities 0.43 S/m, 0.2 S/m, and 1.79 S/m respectively [87–89]. Source space was represented by 8014 dipoles with fixed orientation and orthogonal to the cortical surface. To avoid the influence of developmental changes and inter-individual variability in cortical geometry on our results, we opted to project all EEG data onto cortical template of full-term infant which is about in the middle of the studied age range. Forward operator was computed using symmetric boundary element method [90]. Whereas, inverse operator was computed with dynamic statistical parametric mapping approach [91] as it is implemented in Brainstorm [92]. All sources were further collapsed into 58 cortical parcels using infant parcellation scheme [33]. Next, based on their overlap with brain anatomical regions, all parcels were categorized into frontal, central, occipital, and temporal. Cortical signals were computed as the weighted mean of all underlying source signals within the host parcels. To compute functional brain age (see section below), we used broad-band (0.4–40 Hz) cortical signals. However, to study developmental trajectories of O-information measures, we further filtered cortical signals into four frequency bands of interest: delta (1.5–4 Hz), theta (4–8 Hz), alpha (8–13 Hz), and beta (13–22Hz). All band-pass filtering in this work was implemented by using combinations of high-pass and low-pass Butterworth filters with the corresponding cut-offs. Filters were applied offline and in forward-backward directions to avoid distortions of phases caused by infinite impulse response filters. The attenuation in the stopband in one direction was at least 15 dB.

### 4.5 Association with neurodevelopment

We correlated (Spearman test) O-information measures computed for infant brain around term age (38-40 weeks) to Bayley cognitive scores [25] assessed at 1.5 years of age. Following the recommended criteria [93], we excluded subjects with moderate and severe neurodevelopmental delay (cognitive scores *<* 85) from this analysis. For subjects with two recordings (N = 3) falling into the age range of interest, we used average of their O-information indices. Consequently, N = 41 infants were included in the correlation analysis. To exclude the impact of the developmental changes, we also computed same correlations after regressing conceptional ages from O-information measures (see Figure 3A). Finally, we used Benjamini-Hochberg procedure to control for multiple comparisons (across four frequency bands).

### 4.6 Higher-Order Information Analysis

To asses the emergence of higher-order, coordinated brain activity involving multiple regions, we used measures from information theory [94]. When considering how groups of three or more brain regions share information, there is a distinction to be made between different kinds of higher-order interaction [95]. Some information is stored *redundantly*: it is duplicated over individual brain regions and so could be learned by observing Region 1 alone or Region 2 alone or Regions 3 alone, and so on. The alternative is information that is stored *synergistically*, in the joint-state of two or more regions. This is information that can only be learned by knowing the state of Region 1 and Region 2 and Region 3, and so on. For a more detailed discussion of redundancy, synergy, and logical implicature, see [96, 97]. Synergistic information requires a high degree of coordination between multiple regions, forming an integrated “whole” that is “greater than the sum of it’s parts” [19].

To explore the distribution and redundancies across the developing neonatal cortex, we used a recently proposed, information-theoretic measure: the O-information [20]. A heuristic measure, for a multivariate system **X**, the O-information of that system, Ω(**X**) reflects whether the information structure of the system is redundancy-dominated (in which case, Ω(**X**) *>* 0) or synergy-dominated (in which case, Ω(**X**) *<* 0.

For a more detailed, mathematical analysis of the O-information, see the original proposal by Rosas et al., [20]. Briefly, the O-information begins with a simpler measure that generalizes the bivariate Shannon mutual information to arbitrarilly large systems. Originally introduced by Watanabe as the total correlation [98] and then independently re-derived by Tononi, Sporns, and Edelman as the integration [99], the total correlation is defined by:

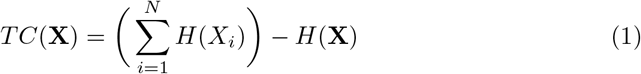

where *N* = |**X**|, and *H*() is the Shannon entropy function. The total correlation can be thought of as a measure of redundancy: *TC*(**X**) is maximal when every *X*_*i*_ is a copy of every other variable. Varley et al., showed that the O-information can be written in terms of sums and differences of total correlations [19]:

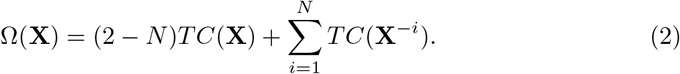

Where **X**^*−i*^ indicates the joint state of every element of **X** excluding *X*_*i*_. For instance, if **X** = *{X*_1_, *X*_2_, *X*_3_, *X*_4_*}*, then **X**^*−*2^ = *{X*_1_, *X*_3_, *X*_4_*}*. We can intuitively understand Ω(**X**) as quantifying the difference between the integration of the “whole” and the integration the “parts.” The left-hand term, (2 *− N*)*TC*(**X**) is the integration of the whole **X**, duplicated (2 *− N*) times (and is therefore a large, negative number, as *N* is always greater than two in higher-order interactions). The righthand side, 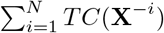can be understood as adding back in the integration of every lower-order ensemble that excludes one element each time. If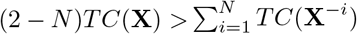, then there is integration in the whole that is not accounted for by the sum of the lower order parts. This is why Ω(**X**) *<* 0 is taken as a heuristic indicator of synergy.

Given the continuous nature of electrophysiological signals, we used Gaussian estimators of the differential total correlation [99]:

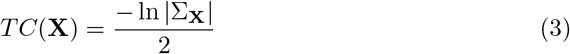

where |Σ_**X**_| is the determinant of the covariance matrix of **X**. For every recording, we computed global O-information and global total correlation for the entire brain.

### 4.7 Simulated Annealing to Extract Synergistic Subsystems

Following Varley et al., [19], we used a variant of the simulated annealing algorithm [23] to extract maximally synergistic ensembles of brain regions. Briefly, given a fixed ensemble size of *k*, the simulated annealing algorithm begins with a random set of *k* brain regions, and then swaps regions in and out to minimize the objective function (in this case, the O-information). Prior work in adult, human neuroimaging data has shown that optimally synergistic O-information has a non-monotonic relationship with *k* [19]. Consequently, we ran the optimization for every integer value of *k* in the range 3-50.

Since O-information is a measure of redundancy/synergy bias, for a given optimal ensemble of size *k*, it cannot be assumed that the *k* elements are not “compromised” by the presence of redundancy. For example, one could imagine that the annealing algorithm returns a set **X**^*∗*^ of five regions, where three regions are highly synergistic amongst themselves, while the other two regions are independent, or perhaps weakly redundant. Claiming that the Ω(**X**^*∗*^) represents a synergistic dependency between all five elements would be spurious. Following [19], for every optimal bag **X**^*∗*^, we used a filtering algorithm: if the removal of any single element from **X**^*∗*^ decreases the Oinformation, then we say that the set **X**^*∗*^ is not “irreducibly synergistic” and removed it from our analysis.

### 4.8 Developmental trajectories

We produced developmental trajectories for infants’ subgroups (FNI and SC) by computing mean values within two-weeks-wide sliding windows with 50% overlap (Fig.4A). These settings were selected as a compromise between temporal resolution and the sample sizes. Further, clinical groups were compared in each age bin using Wilcoxon rank-sum test. For the age bins which showed consistent group differences in Oinformation across all frequency bands (35 and 38 weeks), we also compared minima of O-information and optimal subsystem size. The same approach was used also for functional brain age courses (Fig.4D). Effect size of the group differences was estimated using rank-biserial correlation (*r*).

### 4.9 Computation of functional brain age

Functional brain age (FBA) was estimated based on the combination of several features extracted from the 5-minute-long cortical signals. A total of N = 43 features extracted from the EEG for each epoch (see https://github.com/nstevensonUH/Neonatal-EEG-Analysis for a public repository of Matlab code to calculate features). These features were designed to summarise the amplitude, frequency, and information content of the brain signals; cross-channel/parcel measurements were removed as regionally specific evaluation of FBA was the aim of analysis [49]. Features were estimated from each cortical signal and then averaged across all parcels within an anatomical region (four regions were used: frontal, central, temporal, and occipital). The 43 features per region were then combined using support vector regression to calculate FBA estimate [100]. The combination was trained within a 10-fold cross validation where approximately 90% of cortical signals were included in a training set and 10% of cortical signals were left out for testing; a process that was repeated 10 times until all data had been tested on (due to the presence of multiple EEG recordings and twins in the dataset, cross-validation selections were based on mother’s ID number). Within each training fold, feature selection was applied using a hybrid filter-wrapper approach to reduce the dimensionality of the input feature vector [101]. As a first stage, only features with a significant correlation (corrected for multiple comparisons; Benjamini–Hochberg procedure) with age were selected (filter stage). The residual feature set was applied to a backwards feature selection as a second stage, with a stopping criterion based on the Akaike Information Criterion evaluated on an internal 4-fold cross-validation (wrapper stage). The SVR was trained using the Matlab function fitrsvm.m with a medium Gaussian kernel (Kernel Scale = 9.8, box constraint = IQR/1.349, *ϵ* = IQR/13.49 and IQR is the interquartile range the input PMA). The process of training the FBA was for each anatomical region resulting in 4 FBAs per infant EEG recording.

## Declarations

## Acknowledgements

This work was funded by The Finnish Academy (335778, 332017 to S.V. and 321235 to A.T.), the Jusélius Foundation (S.V., A.T.). T.F.V. was supported by NSF-NRT grant 1735095, Interdisciplinary Training in Complex Networks and Systems. T.F.V. would like to thank Mrs. Maria Pope for her expertise on simulated annealing. We extend our gratitude to Dr. Pauliina Yrjölä for her assistance in EEG data preparation.

## Conflict of interest statement

The authors declare no competing interests.

## Notes

### Competing Interest Statement

The authors have declared no competing interest.

## References

[1] Gutknecht, A.J., Wibral, M., Makkeh, A.: Bits and pieces: understanding information decomposition from part-whole relationships and formal logic. Proc Math Phys Eng Sci 477(2251), 20210110 (2021)10.1098/rspa.2021.0110

[2] Luppi, A.I., Mediano, P.A.M., Rosas, F.E., Holland, N., Fryer, T.D., O’Brien, J.T., Rowe, J.B., Menon, D.K., Bor, D., Stamatakis, E.A.: A synergistic core for human brain evolution and cognition. Nat Neurosci 25(6), 771–782 (2022) 10.1038/s41593-022-01070-0

[3] Luppi, A.I., Rosas, F.E., Mediano, P.A.M., Menon, D.K., Stamatakis, E.A.: Information decomposition and the informational architecture of the brain. Trends Cogn Sci (2024) 10.1016/j.tics.2023.11.005

[4] Varley, T.F.: Information theory for complex systems scientists. ArXiv abs/2304.12482 (2023)

[5] Gatica, M., Cofré, R., Mediano, P.A.M., Rosas, F.E., Orio, P., Diez, I., Swinnen, S.P., Cortes, J.M.: High-order interdependencies in the aging brain. Brain Connectivity 11(9), 734–744 (2021) 10.1089/brain.2020.0982

[6] Newman, E.L., Varley, T.F., Parakkattu, V.K., Sherrill, S.P., Beggs, J.M.: Revealing the dynamics of neural information processing with multivariate information decomposition. Entropy 24(7), 930 (2022)

[7] Varley, T.F., Pope, M., Maria, G., Joshua, Sporns O.: Partial entropy decomposition reveals higher-order information structures in human brain activity. Proc Natl Acad Sci U S A 120(30), 2300888120 (2023)10.1073/pnas.2300888120

[8] Varley, T.F., Sporns, O., Schaffelhofer, S., Scherberger, H., Dann, B.: Information-processing dynamics in neural networks of macaque cerebral cortex reflect cognitive state and behavior. Proceedings of the National Academy of Sciences 120(2), 2207677120 (2023) 10.1073/pnas.2207677120

[9] Tokariev, A., Videman, M., Palva, J.M., Vanhatalo, S.: Functional brain connectivity develops rapidly around term age and changes between vigilance states in the human newborn. Cereb Cortex 26(12), 4540–4550 (2016) 10.1093/cercor/bhv219

[10] Batalle, D., Hughes, E.J., Zhang, H., Tournier, J.D., Tusor, N., Aljabar, P., Wali, L., Alexander, D.C., Hajnal, J.V., Nosarti, C., Edwards, A.D., Counsell, S.J.: Early development of structural networks and the impact of prematurity on brain connectivity. Neuroimage 149, 379–392 (2017)10.1016/j.neuroimage.2017.01.065

[11] Keunen, K., Counsell, S.J., Benders, M.J.: The emergence of functional architecture during early brain development. Neuroimage (2017)10.1016/j.neuroimage.2017.01.047

[12] Turk, E., Heuvel, M.I., Benders, M.J., Heus, R., Franx, A., Manning, J.H., Hect, J.L., Hernandez-Andrade, E., Hassan, S.S., Romero, R., Kahn, R.S., Thomason, M.E., Heuvel, M.P.: Functional connectome of the fetal brain. J Neurosci 39(49), 9716–9724 (2019) 10.1523/jneurosci.2891-18.2019

[13] Kostovic, I., Radoš, M., Kostovic-Srzentic, M., Krsnik, Z.: Fundamentals of the development of connectivity in the human fetal brain in late gestation: From 24 weeks gestational age to term. Journal of neuropathology and experimental neurology 80(5), 393–414 (2021) 10.1093/jnen/nlab024

[14] Karolis, V.R., Fitzgibbon, S.P., Cordero-Grande, L., Farahibozorg, S.-R., Price, A.N., Hughes, E.J., Fetit, A.E., Kyriakopoulou, V., Pietsch, M., Rutherford, M.A., Rueckert, D., Hajnal, J.V., Edwards, A.D., O’Muircheartaigh, J., Duff, E.P., Arichi, T.: Maturational networks of human fetal brain activity reveal emerging connectivity patterns prior to ex-utero exposure. Communications Biology 6(1), 661 (2023) 10.1038/s42003-023-04969-x

[15] Luhmann, H.J., Sinning, A., Yang, J.W., Reyes-Puerta, V., Stüttgen, M.C., Kirischuk, S., Kilb, W.: Spontaneous neuronal activity in developing neocortical networks: From single cells to large-scale interactions. Front Neural Circuits 10, 40 (2016) 10.3389/fncir.2016.00040

[16] Molnár, Z., Luhmann, H.J., Kanold, P.O.: Transient cortical circuits match spontaneous and sensory-driven activity during development. Science 370(6514) (2020) 10.1126/science.abb2153

[17] Martini, F.J., Guillamón-Vivancos, T., Moreno-Juan, V., Valdeolmillos, M., López-Bendito, G.: Spontaneous activity in developing thalamic and cortical sensory networks. Neuron 109(16), 2519–2534 (2021)10.1016/j.neuron.2021.06.026

[18] Luhmann, H.J., Kanold, P.O., Molnár, Z., Vanhatalo, S.: Early brain activity: Translations between bedside and laboratory. Prog Neurobiol 213, 102268 (2022) 10.1016/j.pneurobio.2022.102268

[19] Varley, T.F., Pope, M., Faskowitz, J., Sporns, O.: Multivariate information theory uncovers synergistic subsystems of the human cerebral cortex. Communications Biology 6(1), 451 (2023) 10.1038/s42003-023-04843-w

[20] Rosas, F.E., Mediano, P.A.M., Gastpar, M., Jensen, H.J.: Quantifying high-order interdependencies via multivariate extensions of the mutual information. Physical Review E 100(3), 032305 (2019)10.1103/PhysRevE.100.032305

[21] Myers, M.M., Grieve, P.G., Stark, R.I., Isler, J.R., Hofer, M.A., Yang, J., Ludwig, R.J., Welch, M.G.: Family Nurture Intervention in preterm infants alters frontal cortical functional connectivity assessed by EEG coherence. Acta Paediatrica (Oslo, Norway: 1992) 104(7), 670–677 (2015)10.1111/apa.13007

[22] Yrjölä, P., Myers, M.M., Welch, M.G., Stevenson, N.J., Tokariev, A., Vanhatalo, S.: Facilitating early parent-infant emotional connection improves cortical networks in preterm infants. Science Translational Medicine 14(664), 4786 (2022) 10.1126/scitranslmed.abq4786

[23] Metropolis, N., Rosenbluth, A.W., Rosenbluth, M.N., Teller, A.H., Teller, E.: Equation of State Calculations by Fast Computing Machines. The Journal of Chemical Physics 21(6), 1087–1092 (1953) 10.1063/1.1699114. Publisher: American Institute of Physics. Accessed 2022-06-08

[24] Xia, M., Wang, J., He, Y.: BrainNet Viewer: A Network Visualization Tool for Human Brain Connectomics. PLOS ONE 8(7), 68910 (2013) 10.1371/journal.pone.0068910. Publisher: Public Library of Science. Accessed 2024-02-23

[25] Bayley, N.: Bayley Scales of Infant and Toddler Development, 3rd. ed edn., p. 1. Harcourt Assessment, San Antonio, TX (2006)

[26] Welch, M.G., Firestein, M.R., Austin, J., Hane, A.A., Stark, R.I., Hofer, M.A., Garland, M., Glickstein, S.B., Brunelli, S.A., Ludwig, R.J., Myers, M.M.: Family nurture intervention in the neonatal intensive care unit improves social-relatedness, attention, and neurodevelopment of preterm infants at 18 months in a randomized controlled trial. Journal of Child Psychology and Psychiatry 56(11), 1202–1211 (2015) 10.1111/jcpp.12405

[27] Alex, A.M., Buss, C., Davis, E.P., Campos, G.d.l., Donald, K.A., Fair, D.A., Gaab, N., Gao, W., Gilmore, J.H., Girault, J.B., Grewen, K., Groenewold, N.A., Hankin, B.L., Ipser, J., Kapoor, S., Kim, P., Lin, W., Luo, S., Norton, E.S., O’Connor, T.G., Piven, J., Qiu, A., Rasmussen, J.M., Skeide, M.A., Stein, D.J., Styner, M.A., Thompson, P.M., Wakschlag, L., Knickmeyer, R.: Genetic influences on the developing young brain and risk for neuropsychiatric disorders. Biological Psychiatry 93(10), 905–920 (2023)10.1016/j.biopsych.2023.01.013

[28] Zhou, Y., Song, H., Ming, G.-l.: Genetics of human brain development. Nature Reviews Genetics 25(1), 26–45 (2024)10.1038/s41576-023-00626-5

[29] Sale, A.: A systematic look at environmental modulation and its impact in brain development. Trends in Neurosciences 41(1), 4–17 (2018)10.1016/j.tins.2017.10.004

[30] Miguel, P.M., Pereira, L.O., Silveira, P.P., Meaney, M.J.: Early environmental influences on the development of children’s brain structure and function. Dev Med Child Neurol 61(10), 1127–1133 (2019)10.1111/dmcn.14182

[31] Nolvi, S., Merz, E.C., Kataja, E.-L., Parsons, C.E.: Prenatal stress and the developing brain: Postnatal environments promoting resilience. Biological Psychiatry 93(10), 942–952 (2023) 10.1016/j.biopsych.2022.11.023

[32] Rubinstein, M.R., Burgueño, A.L., Quiroga, S., Wald, M.R., Genaro, A.M.: Current understanding of the roles of gut-brain axis in the cognitive deficits caused by perinatal stress exposure. Cells 12(13) (2023)10.3390/cells12131735

[33] Tokariev, A., Roberts, J.A., Zalesky, A., Zhao, X., Vanhatalo, S., Breakspear, M., Cocchi, L.: Large-scale brain modes reorganize between infant sleep states and carry prognostic information for preterms. Nat Commun 10(1), 2619 (2019) 10.1038/s41467-019-10467-8

[34] Inder, T.E., Volpe, J.J., Anderson, P.J.: Defining the neurologic consequences of preterm birth. N Engl J Med 389(5), 441–453 (2023)10.1056/NEJMra2303347

[35] Russ, J.B., Ostrem, B.E.L.: Acquired brain injuries across the perinatal spectrum: Pathophysiology and emerging therapies. Pediatr Neurol 148, 206–214 (2023) 10.1016/j.pediatrneurol.2023.08.001

[36] Syvälahti, T., Tuiskula, A., Nevalainen, P., Metsäranta, M., Haataja, L., Vanhatalo, S., Tokariev, A.: Networks of cortical activity show graded responses to perinatal asphyxia. Pediatr Res (2023) 10.1038/s41390-023-02978-4

[37] Georgieff, M.K., Ramel, S.E., Cusick, S.E.: Nutritional influences on brain development. Acta Paediatrica 107(8), 1310–1321 (2018)10.1111/apa.14287

[38] Fitzgerald, E., Hor, K., Drake, A.J.: Maternal influences on fetal brain development: The role of nutrition, infection and stress, and the potential for intergenerational consequences. Early Hum Dev 150, 105190 (2020) 10.1016/j.earlhumdev.2020.105190

[39] Tokariev, A., Breakspear, M., Videman, M., Stjerna, S., Scholtens, L.H., Heuvel, M.P., Cocchi, L., Vanhatalo, S.: Impact of in utero exposure to antiepileptic drugs on neonatal brain function. Cereb Cortex 32(11), 2385–2397 (2022) 10.1093/cercor/bhab338

[40] Han, Y., Yuan, M., Guo, Y.S., Shen, X.Y., Gao, Z.K., Bi, X.: The role of enriched environment in neural development and repair. Front Cell Neurosci 16, 890666 (2022) 10.3389/fncel.2022.890666

[41] Lordier, L., Meskaldji, D.-E., Grouiller, F., Pittet, M.P., Vollenweider, A., Vasung, L., Borradori-Tolsa, C., Lazeyras, F., Grandjean, D., Van De Ville, D., Hüppi, P.S.: Music in premature infants enhances high-level cognitive brain networks. Proceedings of the National Academy of Sciences 116(24), 12103–12108 (2019) 10.1073/pnas.1817536116

[42] Haslbeck, F.B., Jakab, A., Held, U., Bassler, D., Bucher, H.-U., Hagmann, C.: Creative music therapy to promote brain function and brain structure in preterm infants: A randomized controlled pilot study. NeuroImage: Clinical 25, 102171 (2020) 10.1016/j.nicl.2020.102171

[43] Charpak, N., Tessier, R., Ruiz, J.G., Uriza, F., Hernandez, J.T., Cortes, D., Montealegre-Pomar, A.: Kangaroo mother care had a protective effect on the volume of brain structures in young adults born preterm. Acta Paediatrica 111(5), 1004–1014 (2022) 10.1111/apa.16265

[44] Beltrán, M.I., Dudink, J., Jong, T.M., Benders, M.J.N.L., Hoogen, A.: Sensory-based interventions in the nicu: systematic review of effects on preterm brain development. Pediatric Research 92(1), 47–60 (2022)10.1038/s41390-021-01718-w

[45] Welch, M.G., Hofer, M.A., Brunelli, S.A., Stark, R.I., Andrews, H.F., Austin, J., Myers, M.M.: Family nurture intervention (fni): methods and treatment protocol of a randomized controlled trial in the nicu. BMC Pediatr 12, 14 (2012) 10.1186/1471-2431-12-14

[46] Moore, E.R., Bergman, N., Anderson, G.C., Medley, N.: Early skin-to-skin contact for mothers and their healthy newborn infants. Cochrane Database Syst Rev 11(11), 003519 (2016) 10.1002/14651858.CD003519.pub4

[47] Welch, M.G., Myers, M.M.: Advances in family-based interventions in the neonatal icu. Curr Opin Pediatr 28(2), 163–9 (2016)10.1097/MOP.0000000000000322

[48] Welch, M.G., Grieve, P.G., Stark, R.I., Isler, J.R., Ludwig, R.J., Hane, A.A., Gong, A., Darilek, U., Austin, J., Myers, M.M.: Family nurture intervention increases term age forebrain eeg activity: A multicenter replication trial. Clinical Neurophysiology 138, 52–60 (2022)10.1016/j.clinph.2022.02.018

[49] Stevenson, N.J., Oberdorfer, L., Tataranno, M.L., Breakspear, M., Colditz, P.B., Vries, L.S., Benders, M., Klebermass-Schrehof, K., Vanhatalo, S., Roberts, J.A.: Automated cot-side tracking of functional brain age in preterm infants. Ann Clin Transl Neurol 7(6), 891–902 (2020) 10.1002/acn3.51043

[50] Proca, A.M., Rosas, F.E., Luppi, A.I., Bor, D., Crosby, M., Mediano, P.A.: Synergistic information supports modality integration and flexible learning in neural networks solving multiple tasks. arXiv preprint 2210.02996 (2022)

[51] Vanderhaeghen, P., Polleux, F.: Developmental mechanisms underlying the evolution of human cortical circuits. Nature Reviews Neuroscience 24(4), 213–232 (2023) 10.1038/s41583-023-00675-z

[52] Loomba, S., Straehle, J., Gangadharan, V., Heike, N., Khalifa, A., Motta, A., Ju, N., Sievers, M., Gempt, J., Meyer, H.S., Helmstaedter, M.: Connectomic comparison of mouse and human cortex. Science 377(6602), 0924 (2022) 10.1126/science.abo0924

[53] Schmidt, E.R.E., Zhao, H.T., Park, J.M., Dipoppa, M., Monsalve-Mercado, M.M., Dahan, J.B., Rodgers, C.C., Lejeune, A., Hillman, E.M.C., Miller, K.D., Bruno, R.M., Polleux, F.: A human-specific modifier of cortical connectivity and circuit function. Nature 599(7886), 640–644 (2021)10.1038/s41586-021-04039-4

[54] Knoblauch, K., Ercsey-Ravasz, M., Kennedy, H., Toroczkai, Z.: In: Kennedy, H., Van Essen, D.C., Christen, Y. (eds.) The Brain in Space, pp. 45–74. Springer, Cham (CH) (2016).10.1007/978-3-319-27777-65

[55] Puxeddu, M.G., Faskowitz, J., Seguin, C., Yovel, Y., Assaf, Y., Betzel, R., Sporns, O.: Relation of connectome topology to brain volume across 103 mammalian species. PLoS Biol 22(2), 3002489 (2024)10.1371/journal.pbio.3002489

[56] Van Essen, D.C., Donahue, C.J., Coalson, T.S., Kennedy, H., Hayashi, T., Glasser, M.F.: Cerebral cortical folding, parcellation, and connectivity in humans, nonhuman primates, and mice. Proceedings of the National Academy of Sciences 116(52), 26173–26180 (2019)10.1073/pnas.1902299116

[57] Markov, N.T., Ercsey-Ravasz, M.M., Ribeiro Gomes, A.R., Lamy, C., Magrou, L., Vezoli, J., Misery, P., Falchier, A., Quilodran, R., Gariel, M.A., Sallet, J., Gamanut, R., Huissoud, C., Clavagnier, S., Giroud, P., Sappey-Marinier, D., Barone, P., Dehay, C., Toroczkai, Z., Knoblauch, K., Van Essen, D.C., Kennedy, H.: A weighted and directed interareal connectivity matrix for macaque cerebral cortex. Cereb Cortex 24(1), 17–36 (2014)10.1093/cercor/bhs270

[58] Oh, S.W., Harris, J.A., Ng, L., Winslow, B., Cain, N., Mihalas, S., Wang, Q., Lau, C., Kuan, L., Henry, A.M., Mortrud, M.T., Ouellette, B., Nguyen, T.N., Sorensen, S.A., Slaughterbeck, C.R., Wakeman, W., Li, Y., Feng, D., Ho, A., Nicholas, E., Hirokawa, K.E., Bohn, P., Joines, K.M., Peng, H., Hawrylycz, M.J., Phillips, J.W., Hohmann, J.G., Wohnoutka, P., Gerfen, C.R., Koch, C., Bernard, A., Dang, C., Jones, A.R., Zeng, H.: A mesoscale connectome of the mouse brain. Nature 508(7495), 207–214 (2014) 10.1038/nature13186

[59] Buckner, R.L., Krienen, F.M.: The evolution of distributed association networks in the human brain. Trends in Cognitive Sciences 17(12), 648–665 (2013) 10.1016/j.tics.2013.09.017

[60] Petanjek, Z., Judaš, M., Šimic, G., Rasin, M.R., Uylings, H.B., Rakic, P., Kostovic, I.: Extraordinary neoteny of synaptic spines in the human prefrontal cortex. Proc Natl Acad Sci U S A 108(32), 13281–6 (2011)10.1073/pnas.1105108108

[61] Kolb, B., Mychasiuk, R., Muhammad, A., Li, Y., Frost, D.O., Gibb, R.: Experience and the developing prefrontal cortex. Proc Natl Acad Sci U S A 109 Suppl 2(Suppl 2), 17186–93 (2012) 10.1073/pnas.1121251109

[62] Chini, M., Hanganu-Opatz, I.L.: Prefrontal cortex development in health and disease: Lessons from rodents and humans. Trends Neurosci 44(3), 227–240 (2021) 10.1016/j.tins.2020.10.017

[63] Kolk, S.M., Rakic, P.: Development of prefrontal cortex. Neuropsychopharmacology 47(1), 41–57 (2022) 10.1038/s41386-021-01137-9

[64] Larivière, S., Wael, R., Hong, S.-J., Paquola, C., Tavakol, S., Lowe, A.J., Schrader, D.V., Bernhardt, B.C.: Multiscale structure–function gradients in the neonatal connectome. Cerebral Cortex 30(1), 47–58 (2019)10.1093/cercor/bhz069

[65] Ball, G., Seidlitz, J., O’Muircheartaigh, J., Dimitrova, R., Fenchel, D., Makropoulos, A., Christiaens, D., Schuh, A., Passerat-Palmbach, J., Hutter, J., Cordero-Grande, L., Hughes, E., Price, A., Hajnal, J.V., Rueckert, D., Robinson, E.C., Edwards, A.D.: Cortical morphology at birth reflects spatiotemporal patterns of gene expression in the fetal human brain. PLoS Biol 18(11), 3000976 (2020) 10.1371/journal.pbio.3000976

[66] Nielsen, A.N., Kaplan, S., Meyer, D., Alexopoulos, D., Kenley, J.K., Smyser, T.A., Wakschlag, L.S., Norton, E.S., Raghuraman, N., Warner, B.B., Shimony, J.S., Luby, J.L., Neil, J.J., Petersen, S.E., Barch, D.M., Rogers, C.E., Sylvester, C.M., Smyser, C.D.: Maturation of large-scale brain systems over the first month of life. Cerebral Cortex 33(6), 2788–2803 (2022)10.1093/cercor/bhac242

[67] Larsen, B., Sydnor, V.J., Keller, A.S., Yeo, B.T.T., Satterthwaite, T.D.: A critical period plasticity framework for the sensorimotor-association axis of cortical neurodevelopment. Trends Neurosci 46(10), 847–862 (2023)10.1016/j.tins.2023.07.007

[68] Kostovic, I., Judaš, M.: Prolonged coexistence of transient and permanent circuitry elements in the developing cerebral cortex of fetuses and preterm infants. Dev Med Child Neurol 48(5), 388–93 (2006)10.1017/S0012162206000831

[69] Linker, S.B., Narvaiza, I., Hsu, J.Y., Wang, M., Qiu, F., Mendes, A.P.D., Oefner, R., Kottilil, K., Sharma, A., Randolph-Moore, L., Mejia, E., Santos, R., Marchetto, M.C., Gage, F.H.: Human-specific regulation of neural maturation identified by cross-primate transcriptomics. Curr Biol 32(22), 4797–48075 (2022) 10.1016/j.cub.2022.09.028

[70] Petanjek, Z., Banovac, I., Sedmak, D., Hladnik, A.: Dendritic spines: Synaptogenesis and synaptic pruning for the developmental organization of brain circuits. Adv Neurobiol 34, 143–221 (2023)10.1007/978-3-031-36159-34

[71] Wang, L., Pang, K., Zhou, L., Cebrián-Silla, A., González-Granero, S., Wang, S., Bi, Q., White, M.L., Ho, B., Li, J., Li, T., Perez, Y., Huang, E.J., Winkler, E.A., Paredes, M.F., Kovner, R., Sestan, N., Pollen, A.A., Liu, P., Li, J., Piao, X., García-Verdugo, J.M., Alvarez-Buylla, A., Liu, Z., Kriegstein, A.R.: A crossspecies proteomic map reveals neoteny of human synapse development. Nature 622(7981), 112–119 (2023) 10.1038/s41586-023-06542-2

[72] Padilla, N., Saenger, V.M., Hartevelt, T.J., Fernandes, H.M., Lennartsson, F., Andersson, J.L.R., Kringelbach, M., Deco, G., Åden, U.: Breakdown of whole-brain dynamics in preterm-born children. Cereb Cortex 30(3), 1159–1170 (2020) 10.1093/cercor/bhz156

[73] Wallois, F., Routier, L., Heberlé, C., Mahmoudzadeh, M., Bourel-Ponchel, E., Moghimi, S.: Back to basics: the neuronal substrates and mechanisms that underlie the electroencephalogram in premature neonates. Neurophysiol Clin 51(1), 5–33 (2021) 10.1016/j.neucli.2020.10.006

[74] Yrjölä, P., Stjerna, S., Palva, J.M., Vanhatalo, S., Tokariev, A.: Phase-based cortical synchrony is affected by prematurity. Cereb Cortex 32(10), 2265–2276 (2022) 10.1093/cercor/bhab357

[75] França, L.G.S., Ciarrusta, J., Gale-Grant, O., Fenn-Moltu, S., Fitzgibbon, S., Chew, A., Falconer, S., Dimitrova, R., Cordero-Grande, L., Price, A.N., Hughes, E., O’Muircheartaigh, J., Duff, E., Tuulari, J.J., Deco, G., Counsell, S.J., Hajnal, J.V., Nosarti, C., Arichi, T., Edwards, A.D., McAlonan, G., Batalle, D.: Neonatal brain dynamic functional connectivity in term and preterm infants and its association with early childhood neurodevelopment. Nature Communications 15(1), 16 (2024) 10.1038/s41467-023-44050-z

[76] Noori, R., Park, D., Griffiths, J.D., Bells, S., Frankland, P.W., Mabbott, D., Lefebvre, J.: Activity-dependent myelination: A glial mechanism of oscillatory self-organization in large-scale brain networks. Proc Natl Acad Sci U S A 117(24), 13227–13237 (2020) 10.1073/pnas.1916646117

[77] Faria, J. O., Pivonkova, H., Varga, B., Timmler, S., Evans, K.A., Káradóttir, R.T.: Periods of synchronized myelin changes shape brain function and plasticity. Nat Neurosci 24(11), 1508–1521 (2021)10.1038/s41593-021-00917-2

[78] Back, S.A.: White matter injury in the preterm infant: pathology and mechanisms. Acta Neuropathol 134(3), 331–349 (2017)10.1007/s00401-017-1718-6

[79] McNamara, N.B., Miron, V.E.: Microglia in developing white matter and perinatal brain injury. Neurosci Lett 714, 134539 (2020)10.1016/j.neulet.2019.134539

[80] Forbes, T.A., Goldstein, E.Z., Dupree, J.L., Jablonska, B., Scafidi, J., Adams, K.L., Imamura, Y., Hashimoto-Torii, K., Gallo, V.: Environmental enrichment ameliorates perinatal brain injury and promotes functional white matter recovery. Nature Communications 11(1), 964 (2020)10.1038/s41467-020-14762-7

[81] Goldstein, E.Z., Pertsovskaya, V., Forbes, T.A., Dupree, J.L., Gallo, V.: Prolonged environmental enrichment promotes developmental myelination. Front Cell Dev Biol 9, 665409 (2021) 10.3389/fcell.2021.665409

[82] DeMaster, D., Bick, J., Johnson, U., Montroy, J.J., Landry, S., Duncan, A.F.: Nurturing the preterm infant brain: leveraging neuroplasticity to improve neurobehavioral outcomes. Pediatr Res 85(2), 166–175 (2019)10.1038/s41390-018-0203-9

[83] Bourel-Ponchel, E., Hasaerts, D., Challamel, M.-J., Lamblin, M.-D.: Behavioral-state development and sleep-state differentiation during early ontogenesis. Neurophysiologie Clinique 51(1), 89–98 (2021)10.1016/j.neucli.2020.10.003

[84] André, M., Lamblin, M.D., d’Allest, A.M., Curzi-Dascalova, L., MoussalliSalefranque, F., S Nguyen The, T., Vecchierini-Blineau, M.F., Wallois, F., Walls-Esquivel, E., Plouin, P.: Electroencephalography in premature and full-term infants. developmental features and glossary. Neurophysiol Clin 40(2), 59–124 (2010) 10.1016/j.neucli.2010.02.002

[85] Vanhatalo, S., Kaila, K.: Development of neonatal eeg activity: from phenomenology to physiology. Semin Fetal Neonatal Med 11(6), 471–8 (2006) 10.1016/j.siny.2006.07.008

[86] Tokariev, A., Stjerna, S., Lano, A., Metsäranta, M., Palva, J.M., Vanhatalo, S.: Preterm birth changes networks of newborn cortical activity. Cereb Cortex 29(2), 814–826 (2019) 10.1093/cercor/bhy012

[87] Despotovic, I., Cherian, P.J., De Vos, M., Hallez, H., Deburchgraeve, W., Govaert, P., Lequin, M., Visser, G.H., Swarte, R.M., Vansteenkiste, E., Van Huffel, S., Philips, W.: Relationship of eeg sources of neonatal seizures to acute perinatal brain lesions seen on mri: a pilot study. Hum Brain Mapp 34(10), 2402–17 (2013) 10.1002/hbm.22076

[88] Odabaee, M., Tokariev, A., Layeghy, S., Mesbah, M., Colditz, P.B., Ramon, C., Vanhatalo, S.: Neonatal eeg at scalp is focal and implies high skull conductivity in realistic neonatal head models. Neuroimage 96, 73–80 (2014) 10.1016/j.neuroimage.2014.04.007

[89] Tokariev, A., Vanhatalo, S., Palva, J.M.: Analysis of infant cortical synchrony is constrained by the number of recording electrodes and the recording montage. Clin Neurophysiol 127(1), 310–23 (2016)10.1016/j.clinph.2015.04.291

[90] Gramfort, A., Papadopoulo, T., Olivi, E., Clerc, M.: Openmeeg: opensource software for quasistatic bioelectromagnetics. Biomed Eng Online 9, 45 (2010) 10.1186/1475-925X-9-45

[91] Dale, A.M., Liu, A.K., Fischl, B.R., Buckner, R.L., Belliveau, J.W., Lewine, J.D., Halgren, E.: Dynamic statistical parametric mapping: combining fmri and meg for high-resolution imaging of cortical activity. Neuron 26(1), 55–67 (2000) 10.1016/s0896-6273(00)81138-1

[92] Tadel, F., Baillet, S., Mosher, J.C., Pantazis, D., Leahy, R.M.: Brainstorm: a user-friendly application for meg/eeg analysis. Comput Intell Neurosci 2011, 879716 (2011) 10.1155/2011/879716

[93] Johnson, S., Moore, T., Marlow, N.: Using the bayley-iii to assess neurodevelopmental delay: which cut-off should be used? Pediatr Res 75(5), 670–4 (2014) 10.1038/pr.2014.10

[94] Cover, T.M., Thomas, J.A.: Elements of Information Theory. John Wiley & Sons, ??? (2012)

[95] Rosas, F.E., Mediano, P.A.M., Luppi, A.I., Varley, T.F., Lizier, J.T., Stramaglia, S., Jensen, H.J., Marinazzo, D.: Disentangling high-order mechanisms and high-order behaviours in complex systems. Nature Physics, 1–2 (2022) 10.1038/s41567-022-01548-5. Publisher: Nature Publishing Group. Accessed 2022-03-22

[96] Makkeh, A., Gutknecht, A.J., Wibral, M.: Introducing a differentiable measure of pointwise shared information. Physical Review E 103(3), 032149 (2021) 10.1103/PhysRevE.103.032149. Publisher: American Physical Society. Accessed 2021-05-20

[97] Varley, T.F., Kaminski, P.: Untangling Synergistic Effects of Intersecting Social Identities with Partial Information Decomposition. Entropy 24(10), 1387 (2022) 10.3390/e24101387. Number: 10 Publisher: Multidisciplinary Digital Publishing Institute. Accessed 2022-09-28

[98] Watanabe, S.: Information Theoretical Analysis of Multivariate Correlation. IBM Journal of Research and Development 4(1), 66–82 (1960) 10.1147/rd.41.0066. Number: 1

[99] Tononi, G., Sporns, O., Edelman, G.M.: A measure for brain complexity: relating functional segregation and integration in the nervous system. Proceedings of the National Academy of Sciences 91(11), 5033–5037 (1994)10.1073/pnas.91.11.5033. Number: 11. Accessed 2019-05-15

[100] Smola, A.J., Schölkopf, B.: A tutorial on support vector regression. Statistics and Computing 14(3), 199–222 (2004) 10.1023/B:STCO.0000035301.49549.88

[101] Stevenson, N.J., Nordvik, T., Espeland, C.N., Giordano, V., Moltu, S.J., Larsson, P.G., Klebermaß-Schrehof, K., Stiris, T., Vanhatalo, S.: Inter-site generalizability of eeg based age prediction algorithms in the preterm infant. Physiol Meas 44(7) (2023) 10.1088/1361-6579/ace755

